# Single cell analysis of spondyloarthritis regulatory T cells identifies distinct synovial gene expression patterns and clonal fates

**DOI:** 10.1101/2021.05.31.444674

**Authors:** Davide Simone, Frank Penkava, Anna Ridley, Stephen Sansom, M Hussein Al-Mossawi, Paul Bowness

## Abstract

Regulatory T cells (Tregs) play an important role in controlling inflammation and limiting autoimmunity, but their phenotypes at inflammatory sites in human disease are poorly understood. We here analyze the single-cell transcriptome of >16,000 Tregs obtained from peripheral blood and synovial fluid of two patients with HLA-B27+ ankylosing spondylitis and three patients with psoriatic arthritis, closely related forms of inflammatory spondyloarthritis. We identify multiple Treg clusters with distinct transcriptomic profiles, including, among others, a regulatory CD8^+^ subset expressing cytotoxic markers/genes, and a Th17-like *RORC*+ Treg subset characterized by IL-10 and LAG-3 expression. Synovial Tregs show upregulation of interferon signature and TNF receptor superfamily genes, and marked clonal expansion, consistent with tissue adaptation and antigen contact respectively. Individual synovial Treg clones map to different clusters indicating cell fate divergence. Finally, we demonstrate that LAG-3 directly inhibits IL-12/23 and TNF secretion by patient-derived monocytes, a mechanism with translational potential in SpA. Our detailed characterization of Tregs at an important inflammatory site illustrates the marked specialization of Treg subpopulations.

## Introduction

Regulatory T cells (Tregs) are specialised T lymphocytes that control immune responses during inflammatory and autoimmune processes. Whilst Tregs are characterized by expression of the master transcription factor FOXP3 and the IL-2 receptor alpha chain CD25, they show significant functional heterogeneity and utilize diverse suppressive mechanisms including secretion (or sequestration) of soluble mediators, direct cytotoxicity and contact-dependent receptor inhibition^1^. Integrating environmental signals, they can traffic to specific target organs and adopt organ-specific gene signatures and functions^2^, whilst also maintaining plasticity within tissues^3^. Low dimensional analyses based on phenotypical markers do not fully capture the increasingly apparent functional and transcriptional variety of Tregs, potentially overlooking functional cell states that may play a role in controlling inflammation. The phenotype and transcriptional profile of Tregs is yet to be fully delineated, especially at the single cell level, at many sites of tissue inflammation in humans, including the synovial fluid in the course of inflammatory arthritis, representing an opportunity for the study of local regulatory mechanisms.

The spondyloarthritides (SpA) are a group of chronic immune-mediated arthritic conditions characterized by inflammation of spinal and other joints. The commonest forms of SpA, ankylosing spondylitis (AS) and psoriatic arthritis (PsA), together affect approximately 1% of the population, and are characterized by complex immune dysregulation, largely genetically predisposed but with likely common environmental triggers^4^. While the role of effector immunity, and of type 17 immunity in particular, is widely recognized in SpA^5^, the impact and phenotype of Tregs is largely unknown.

Tregs undergo thymic selection and express a unique rearranged T cell receptor (TCR) alpha-beta chain pair. Although Tregs specific for exogenous antigens have been described^6, 7^, they are thought to recognize self-peptides more frequently than conventional T cells with a resulting skewed TCR repertoire^8^. There is evidence that TCR engagement can shape the gene signature of Tregs^9^, but detailed analysis of Tregs antigen specificity has proven challenging because of their relative rarity. Nevertheless, antigen-specific modulation by Tregs could constitute a potential advancement for cell-based therapy of autoimmune diseases. Thus, a deeper understanding of the role of antigens in human Treg biology is very important.

We here report single cell RNA sequencing of approximately 17,000 Tregs from the blood and inflamed joints of patients with Ankylosing Spondylitis and Psoriatic Arthritis, allowing us to define an atlas of Tregs in the context of active joint inflammation. We identify functionally distinct specialized Treg clusters (some novel) with unique gene expression programs, and describe specific changes in transcriptional profile occurring in synovial fluid (SF) Tregs, providing insight into Treg adaptation during inflammation. Furthermore, pairing gene expression analysis with TCR sequencing, we identify clonally expanded and likely antigen-driven Tregs in the SF, and show for the first time functional heterogeneity within individual Treg clones. Among the specialized Treg subpopulations, we describe two LAG-3-expressing Treg subsets (with coexistent cytotoxic and Th17-like features), and show that LAG-3 can directly control inflammatory responses in myeloid cells from SpA patients.

## Results

### Single cell RNA expression profiling of Tregs from HLA-B27+ Ankylosing Spondylitis synovial fluid and blood reveals diverse Treg clusters

To characterize the transcriptional landscape of Tregs in patients with SpA, we used fluorescent activated cell sorting (FACS) to isolate CD3^+^CD45RA^-^CD25^+^CD127^low^ memory Tregs (see **Methods,** **Fig. 1A**, and **Supplementary Fig. 1A**) from the peripheral blood (PB) and SF of 2 patients with HLA-B27+ Ankylosing Spondylitis presenting with active knee arthritis. Single-cell RNA sequencing (scRNA-seq) including 5’ V(D)J 10x Genomics technology allowed exploration of their immune TCR repertoire together with transcriptional definition. We did not include CD4 in the sorting strategy to allow us to capture all regulatory T cells including previously described CD8^+^ Tregs^10^. After careful quality control to remove doublets and low-quality cells (**Methods**), we obtained 13,397 single-cell Treg transcriptomes from both PB and SF. Through sample integration and unsupervised clustering, we identified 10 Treg clusters (**Fig. 1B**). All clusters were present in both patients (**Supplementary Fig. 1B**) and in both PB and SF. Although none of the clusters were exclusively found in one compartment, SF showed enrichment of Canonical, Cycling, Cytotoxic, CCR4/Helios+ and IFN (interferon) signature clusters. Conversely CCR7+ and KLRB1+ Tregs were enriched in blood (**Fig. 1C**). Notably all clusters expressed the lineage-defining genes *FOXP3* and *IL2RA* at comparable levels (**Fig. 1D**).

**Figure 1.**
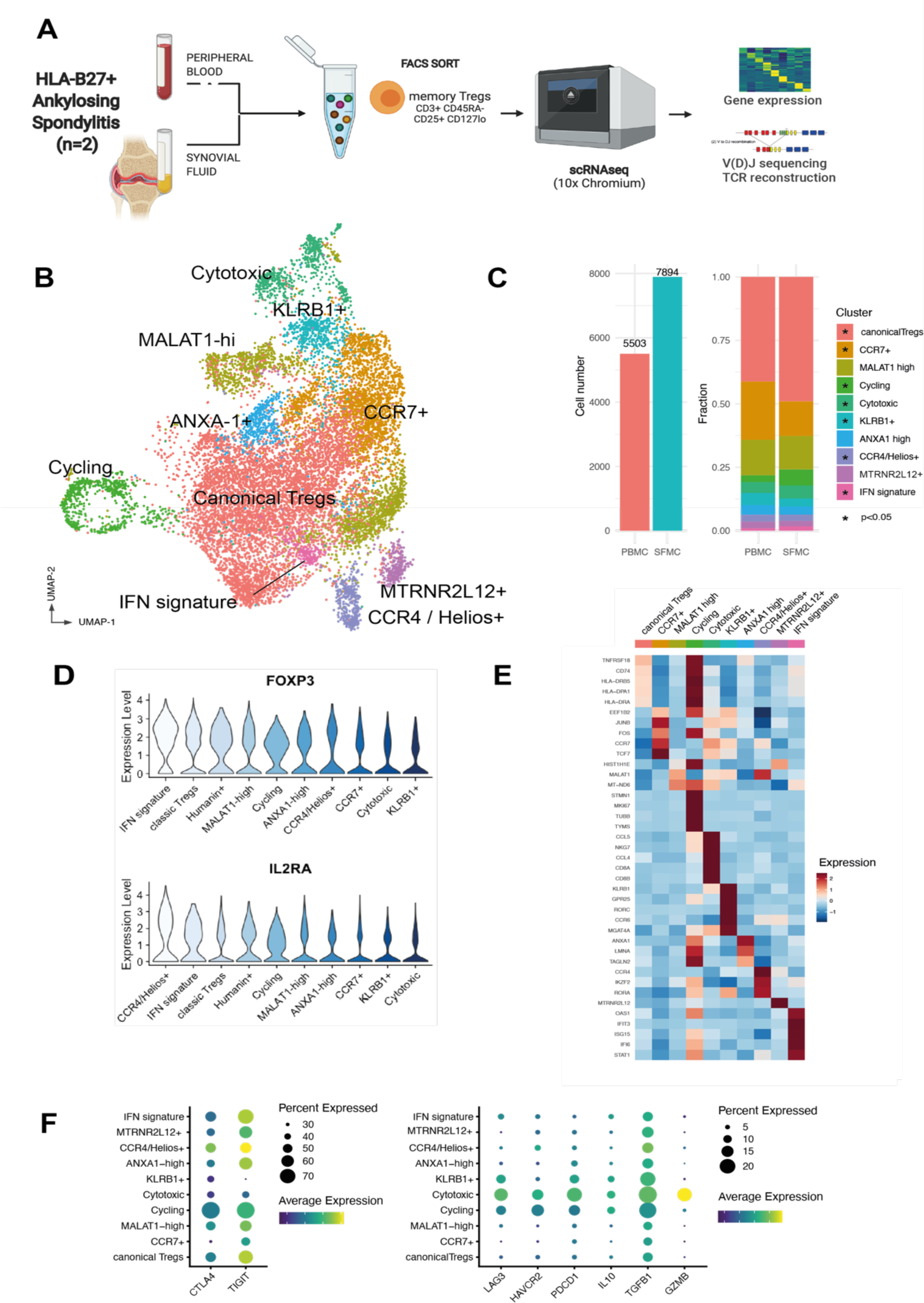
Single cell RNA sequencing analysis of Tregs from HLA-B27+ Ankylosing Spondylitis blood and synovial fluid reveals multiple distinct clusters. (**A**) Experimental design of scRNA-seq of Tregs including gene expression and V(D)J TCR gene segment sequencing. PBMCs and synovial fluid mononuclear cells (SFMCs) from 2 patient with active HLA-B27+ AS were sampled during an arthritis flare. (**B**) Reduced dimensionality visualisation (UMAP plot) and clustering of the transcriptome of 13,397 Tregs. (**C**) Cell numbers from PB and SF and fractions composing each tissue. The asterisks indicate clusters enriched in either PB (CCR7+, KLRB1+) or SF (all the others) (* p<0.05, z-test of two proportions). (**D**) Expression (scaled log(UMI+1)) of lineage defining markers *FOXP3* and *IL2RA* across various clusters, sorted by average expression from the highest-expressing cluster. (E) Heat map of row-wise *z*-score–normalized mean expression of selected marker genes, chosen among top differentially expressed for each cluster and other known markers. (F) Distribution of genes with known T regulatory function across various clusters. Dotplot heatmap showing average scaled expression (color) and percentage of cells (dot size) expressing the genes, split into highly expressed (left panel) and cluster-specific (right panel).

To characterise each cluster and to assist with the annotation, we performed multiple pairwise differential gene expression analyses (**Fig. 1E**, **Supplementary Fig. 1C** and **Supplementary Data 1**). The largest cluster was characterized by high expression levels of canonical Treg genes including *FOXP3*, *TIGIT*, *CD27* and *TNFRSF18*. The second biggest cluster (enriched in blood) expressed high levels of *CCR7* and the transcription factors *JUNB* and *TCF7*. Other distinct cell clusters were characterised by specialised functional and lineage markers (for example *KLRB1*, which encodes CD161, or *GZMA* and *GZMB*, indicative of cytotoxic function) or cell state features (e.g. the cycling cluster or the *MTRNR2L12*+ cluster, whose eponymous marker is a mitochondrial gene). Pathway analysis revealed putative functional pathways for each cluster (**Supplementary Fig. 2A**), including a specific cluster with strong enrichment in genes associated with IFN response.

We next analysed the distribution of effector molecules, including coinhibitory receptors, associated with different mechanisms of suppression across the various clusters. **Figure 1F** left hand panel shows that *CTLA4* and *TIGIT* were highly expressed (up to 70% of cells) in multiple clusters, however *TIGIT* was markedly downregulated in the KLRB1+ cluster relative to other clusters. (log fold change -1.02, p=1.1x10^-577^). Other markers were expressed at lower levels but notably the cytotoxic Treg cluster expressed high levels not only of *GMZB* but also of *PDCD1* and *TGFB1. LAG3 (*and to a lesser extent *IL10)* was preferentially expressed by the cytotoxic, KLRB1+ and cycling clusters (**Fig. 1F**, right hand panel). Spearman pairwise correlation analysis showed co-expression of *LAG3* with *IL10* (**Supplementary Fig. 2C**). Co-expression of *ENTPD1* (CD39, which converts ATP to AMP) was also seen with *CTLA4* with *TIGIT* (**Supplementary Fig. 2C**). Notably, we did not observe significant coexistence of checkpoint inhibitors, as commonly described for tumour infiltrating lymphocytes^11^. We then looked for a co-expression network^9^ in our Treg dataset, finding the tightest co-regulated gene pairs were *FOXP3* and *IL2RA*, and *TNFRSF18* and *TNFRSF4* (**Supplementary Fig. 2D)**. Overall, our data clearly demonstrate the existence of multiple Treg populations with distinct phenotype at a major site of tissue inflammation, with specific population enrichment and gene expression patterns.

### Coordinated gene expression patterns characterize Th17-like and cytotoxic Treg subsets in Ankylosing Spondylitis joints

One cluster, which we designated “KLRB1+”, expressed not only *KLRB1* (coding for CD161, a C-type lectin-like receptor, associated with Th17, MAIT and NK cells^12^), but also the Th17 transcription factor RORC (**Fig. 2A**), a broad Th17 gene module (**Fig. 2B**), and *GPR25*, encoding an orphan G protein coupled receptor previously associated with AS^13^. Interestingly, this cluster had lower expression of *TIGIT* and *IKZF2* (the gene encoding the transcription factor Helios). We confirmed that these cells were mostly Helios^-^ and TIGIT^low^ (**Fig. 2C** and **Supplementary Fig. 3A**) by flow cytometry of blood samples from 14 SpA patients. This population shares features with a ROR*γ*t^+^ Helios^-^ Treg subpopulation described in the mouse intestine^14^.

**Figure 2.**
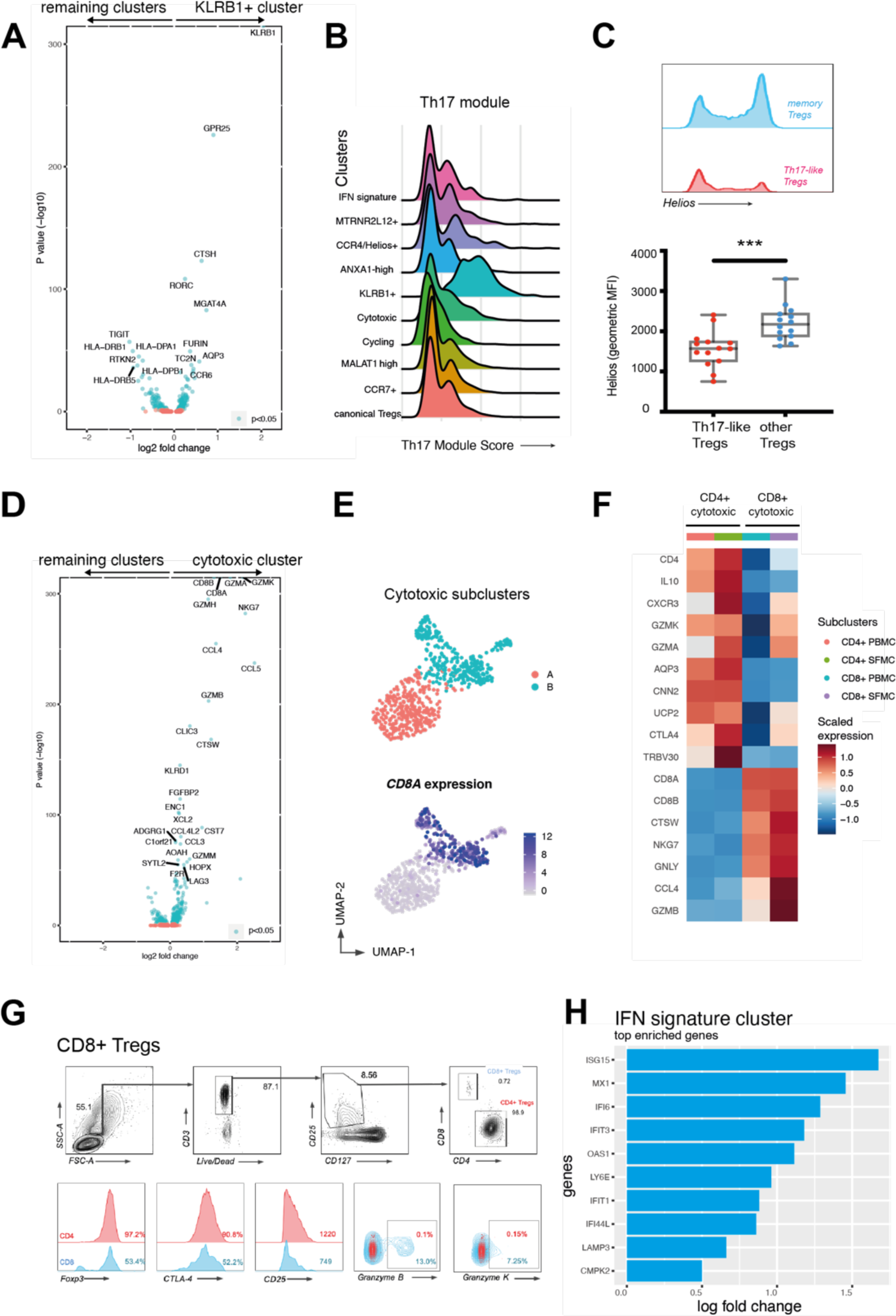
Ankylosing Spondylitis Treg subsets include a Th17-like subset and a cytotoxic subset that contains a CD8-expressing regulatory phenotype. **(A)** Volcano plot shows differential gene expression of KLRB1+ Tregs compared to all other memory Tregs. **(B)** KLRB1+ cluster highly expresses a Th17-like gene module. Distribution of gene score obtained from a curated Th17 gene list (see **Supplementary Table 4**) across the various clusters in the form of ridge plot. **(C)** Representative FACS plot showing Helios expression from Th17-like memory Tregs (CD25^+^CD127^low^CD45RA^-^CCR6^+^) and remaining memory Tregs from an AS patient blood, and Helios expression (geometric mean of fluorescence intensity) in Tregs from 14 SpA PBMCs (gated on CD4^+^ CD25^+^ CD127^low^). *** p< 0.001, Mann Whitney test. **(D)** Volcano plot showing differential gene expression of Cytotoxic Tregs compared to all other Treg clusters. **(E)** After reclustering cytotoxic Tregs, two sub clusters appear (top panel). *CD8A* expression shown in bottom panel. **(F)** Heat map of row-wise *z*-score–normalized mean expression of selected marker genes in CD4+ and CD8+ subclusters. **(G)** CD8+ Treg identification and phenotype on flow cytometry: CD8+ Tregs (defined as CD3^+^ CD8^+^ CD4^-^ CD25^+^ CD127^low^). Representative flow cytometry plots of one of n=8 SpA patients. **(H)** Top 10 upregulated genes in IFN signature cluster.

A second distinct effector Treg subset, that we termed “cytotoxic”, expressed multiple genes associated with cytotoxic function, including granzymes A, K, B and H, and *GNLY* (granulysin) (**Fig. 2D**). This cluster was comprised of two subpopulations, largely separated by the expression of *CD4* and *CD8A*/*CD8B* (**Fig. 2E**) and by their distinct effector programs. The CD4^+^ sub-cluster was additionally enriched for *IL10*, *MAF*, and *CTLA4*. The CD8^+^ sub-cluster had a more marked cytotoxic profile that included *NKG7*, *GNLY* and *GZMB* (**Fig. 2F**), with *FOXP3* and *IL2RA* expression comparable to CD4^+^ Tregs (**Supplementary Fig. 3B**). The presence of a CD8^+^ Treg subset expressing Granzyme B and Granzyme K in the PB and in the SF of patients with SpA, representing up to 1.5% of all the CD3^+^CD25^+^CD127^low^ cells, was confirmed by flow cytometry (**Fig. 2G**). A further subset of interest, predominantly seen in SF, expressed a gene signature indicative of exposure and response to type I and type II interferons (**Fig. 2H**).

### Psoriatic Arthritis synovial fluid (and blood) Tregs contain similar subset identities and gene signatures to AS Tregs

To confirm the findings and validate our observations in a second cohort, we carried out a second analysis of Tregs in psoriatic arthritis (PsA), a related SpA condition. We used a scRNAseq dataset previously published by our group^15^ of FACS-sorted memory CD4^+^ (CD3^+^CD45RA^-^CD4^+^) and CD8^+^ (CD3^+^CD45RA^-^CD8^+^) cells isolated from the SF and PB of 3 PsA patients (**Fig. 3A**). Tregs were identified (using unsupervised clustering) in this dataset as a distinct cluster characterized by significant upregulation of *FOXP3, IL2RA* and *IKZF2*. The raw gene expression data matrix of the PsA Treg cluster comprising 3,066 cells (951 from PB and 2,115 from SF) was exported for in depth downstream analysis (**Methods**).

**Figure 3.**
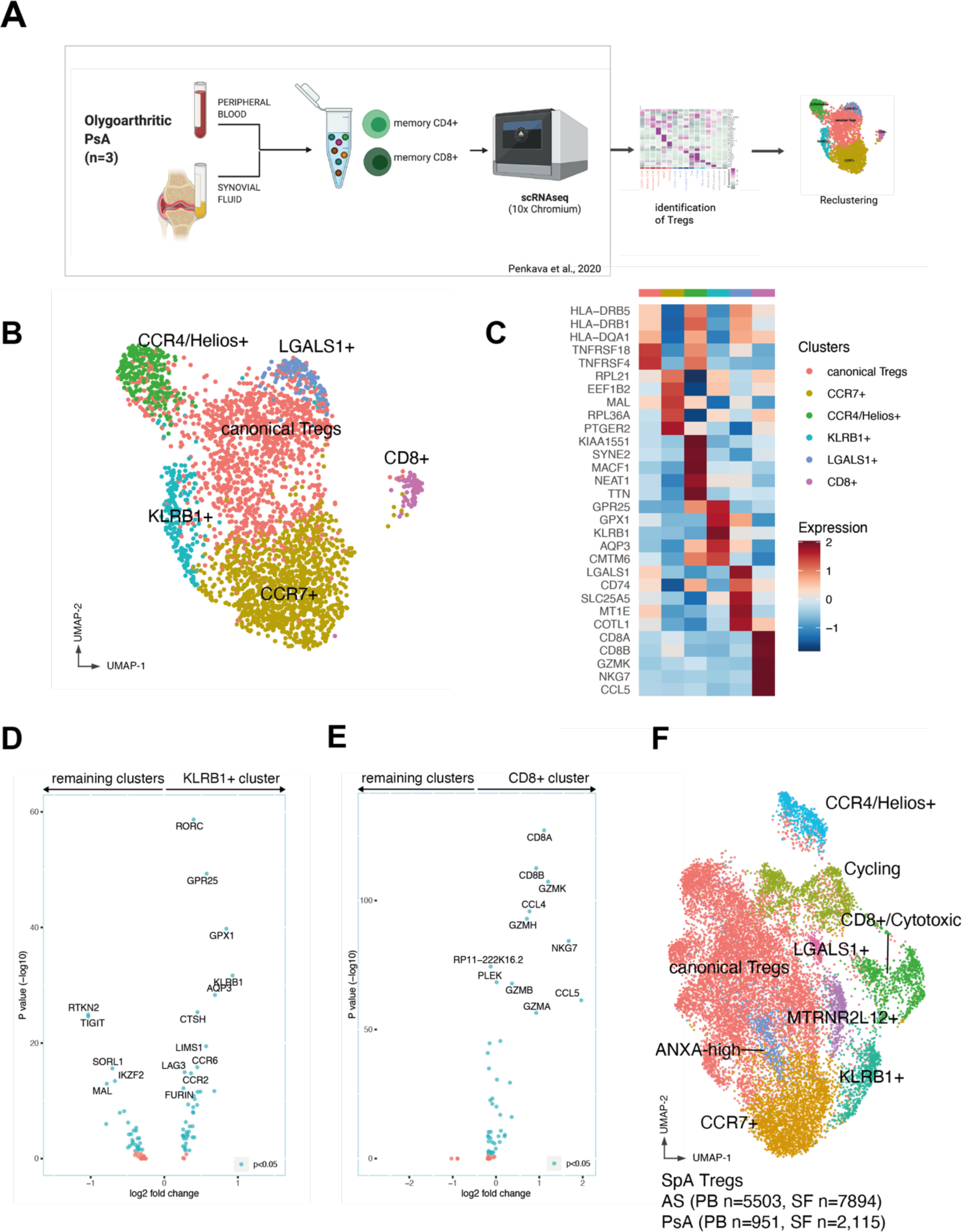
Parallel single cell RNAseq analysis of Psoriatic Arthritis blood and synovial fluid Tregs confirms regulatory subset identities and gene signatures. **(A)** Experimental workflow. The Treg dataset was obtained from ref. 15 (Penkava et al.) identifying the cluster representing Tregs. **(B)** UMAP visualisation of 3,066 Tregs from PB and SF obtained from 3 PsA patients**. (C)** PsA Treg clusters and their differentially expressed genes. Shown is a heatmap of row-wise *z*-score–normalized mean expression of selected marker genes, chosen between top differentially expressed for each cluster. **(D-E)** Differentially expressed genes in KLRB1+ cluster **(D)** and CD8+ cluster **(E)** identified in the PsA dataset compared to the remaining Tregs. **(F)** UMAP plot depicting integrated AS and PsA Treg datasets demonstrating common and overlapping identities.

Six clusters emerged from the analysis, the largest of which expressed conventional Treg markers (*FOXP3, TIGIT* and *TNFRSF18*) and HLA-class II associated genes, very similar to the canonical cluster in the AS dataset (**Fig. 3B-C** and **Supplementary Fig. 4**). Each cluster was found in all three patients and they were similarly distributed in PB and SF (**Supplementary Fig. 4B-C**). Other identified clusters (**Supplementary Data 3**), similarly observed in AS, included a CCR7+ cluster, a CCR4+Helios+ cluster, a *KLRB1*+ cluster, characterised by the expression of *GPR25*, *RORC*, *CCR6*, *IL6R*, *IL1R* and by the downregulation of *TIGIT* and *IKZF2* (**Fig. 3D**) and a CD8+ cytotoxic cluster (**Fig. 3E**). The presence of CD8^+^ Tregs, clustering with the rest of the CD4+ Tregs rather than with CD8^+^ Teffs, suggested that shared transcriptional regulatory signatures prevailed over the expression of lineage markers such as *CD4* and *CD8*. Both the KLRB1+ and cytotoxic clusters from the PsA patients exhibited gene expression profiles closely matching the analogous AS populations described in **Figure 2**, and indeed the two datasets could be readily integrated into a single object including cells with shared transcriptional features from all AS and PsA patients (**Fig. 3F** and **Supplementary Data 4**).

The smaller clusters, indicating rarer phenotype (eg. IFN signature or MTRNR2L12+) that we described in **Fig.1** were not observed in the PsA dataset, potentially due to a smaller sample size. A novel but rare cluster, strongly characterised by expression of *LGALS1* (L-Galectin1), was detected in the synovial fluid and blood from all three PsA patients (**Supplementary Fig. 4D**). A distinct cluster characterised by cell cycle-related genes was not observed, because cycling Tregs clustered together with the other cycling T cells in the original PsA dataset and were thus not found in the downstream analysis.

In summary, Tregs from Psoriatic Arthritis joints and blood closely match those identified in AS, with common subsets and matching gene expression signatures.

### Synovial fluid Tregs upregulate inhibitory markers and show evidence of exposure to TNF and interferons

We next compared the normalised expression of each detected transcript between the SF and PB Tregs in the AS dataset. 401 genes were differentially expressed between the two groups, of which 366 were enriched in SF (**Fig. 4A**). FACS comparison of SpA patient SF and blood confirmed higher cell surface expression of *FOXP3* and CD25 (IL2RA), and also increased numbers of Tregs in the joints (**Supplementary Fig. 5A-B**). The TNF receptor superfamily genes *TNFRSF4* (OX40) and *TNFRSF18* (GITR), and the chemokine *CCL5* (RANTES) were also among the genes upregulated in SF. Annotation of the SF enriched genes using the Reactome resource showed that SF Tregs’ enriched genes were involved in type I and II interferon responses, but also in TCR signalling (**Fig. 4B**). Interestingly, *TNXIP*, encoding a Thioredoxin-interacting protein that regulates redox reactions, was the strongest downregulated gene in the SF. Among the genes upregulated in PB, *CCR6* and *CCR7* were the most striking and consistent with their role in trafficking to the organ and lymphoid structures^16, 17^. Analysis of individual clusters showed frequent recurrence of the same gene pathways (including IFN response pathways) across multiple Treg subsets (**Supplementary Fig. 5C**).

**Figure 4.**
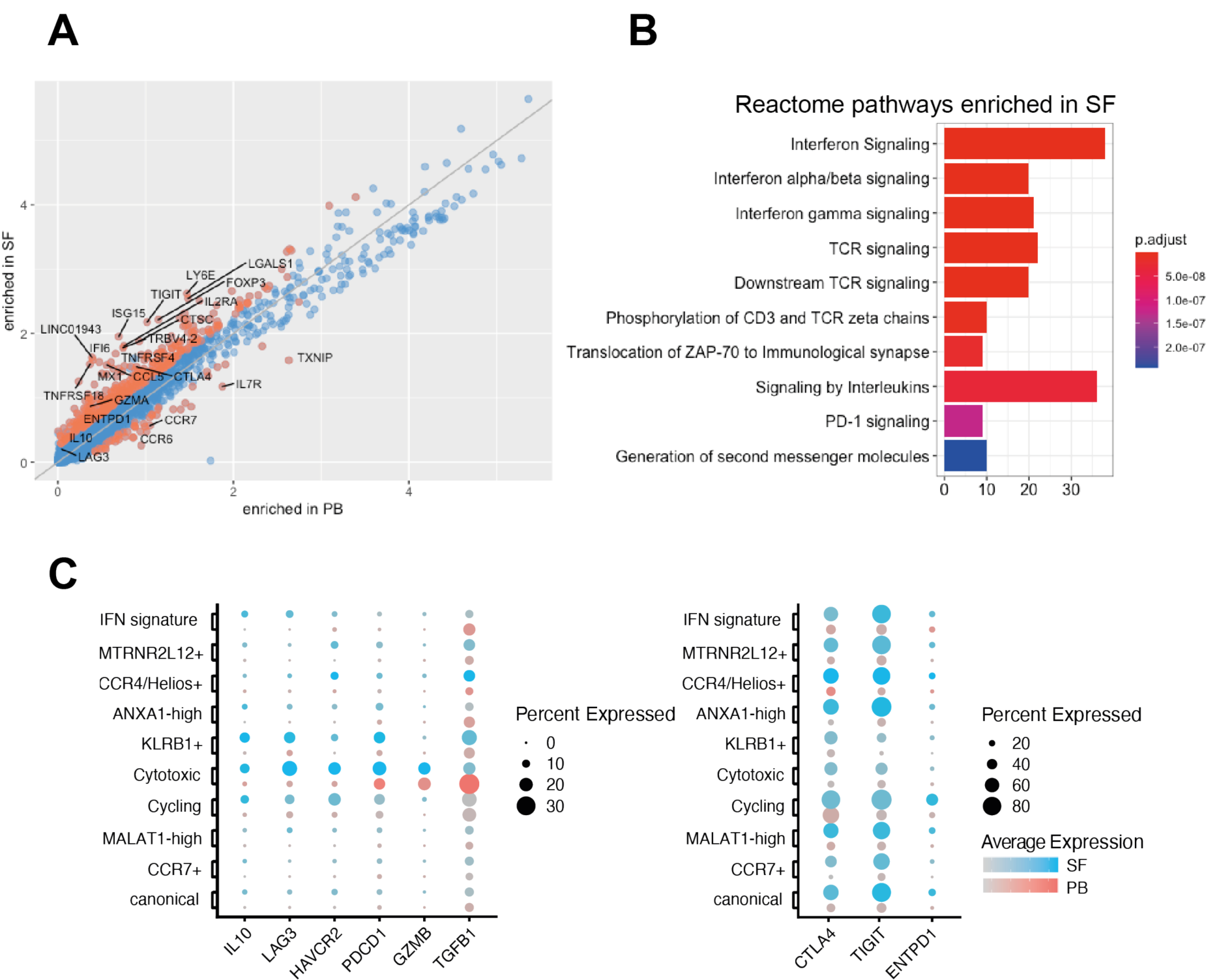
AS Synovial fluid Tregs upregulate inhibitory markers and show evidence of exposure to TNF and interferons. **(A)** Normalised logarithmic (log(UMI+1)) expression of all genes detected in AS PB and SF. Orange dots represent genes that are differentially expressed (Wilcoxon rank-sum test with Bonferroni correction). **(B)** Genes upregulated in AS SF compared to the PB grouped into the top 10 enriched Reactome categories. The color represents the p-value (after Benjamini-Hochberg correction) for each enriched Reactome term. **(C)** Expression of selected suppressive markers in PB (red) and SF (blue dots) clusters. Dots show fraction of expressing cells (dot size), and mean expression level in non-zero cells (colour intensity).

Effector genes were almost always upregulated in the SF (**Fig. 4C**), suggesting an activated phenotype of synovial Tregs, while the expression of genes belonging to the Treg core set was maintained (**Supplementary Fig. 5D-E**), showing that at least at RNA level, these cells remain committed to a suppressive functional programme.

A similar analysis of the PsA dataset (shown in **Supplementary Fig. 5F-G**) revealed very similar genes upregulated in SF, with almost identical pathways observed including most strongly interferon and to a lesser extent TCR signaling.

Thus, both AS and PsA synovial Tregs show transcriptional responses to inflammatory cytokines (particularly IFN, but also TNF), and evidence of TCR signaling (with enhanced expression genes related to suppressive function) consistent with local response to both cytokines and antigen.

### Clonally expanded Tregs with identical TCRα/β sequences and distinct transcriptional features are expanded in SpA (AS and PsA) synovial fluid

The observation of the upregulated TCR signaling pathway genes in synovial Tregs suggested a possible role for cognate antigen in synovial Treg activation. To further explore this, we next determined clonal diversity in our two Treg datasets making use of the novel 5’ chemistry on the 10x Chromium platform to map the TCR *αβ* variable region in our data. Given the large dataset size, we only performed clonal enrichment analysis on clones with at least 3 cells present in either blood or synovial fluid. Cells were defined as belonging to the same clone if they had identical TCR *α* and *β* nucleotide sequences (**Methods**). Based on this assumption we identified 13 statistically enriched clones in AS01 and 52 clones in AS02. The majority of the TCR clonotypes, and the totality of those larger than 100 cells were enriched in the SF (**Fig. 5A**), and exclusively found in the CD4+ compartment. Next, we explored the gene expression of the SF enriched clones compared to the PB enriched clones (**Fig. 5B** and **Supplementary Data 5**). The upregulation of *TIGIT*, *TNFRSF18*, *LGALS1* largely recapitulates the SF signature described in **Figure 4**, with the additional detection of cycling markers (*TUBB*, *TUBA1B*) and *CD177*. Given the prevalent localisation of the Treg clones in the SF rather than the PB, in order to identify the specific transcriptional features of expanded clones, we next compared the differential gene expression of the top 5 SF clones versus the remaining SF cells (**Fig. 5C**). Among the top upregulated genes (apart from TCR *α* and *β* chain variable gene transcripts, not shown), were *CD177* (whose product is the glycosyl-phosphatidylinositol anchored glycoproteins NB1), *SIRPG* (encoding SIRPβ2) and the alarmins *S100A6* and *S100A4* (together with cell cycle markers). Interestingly, *KLRB1*, *CCR6* and *CCR7* were all downregulated in the enriched Treg clones (**Supplementary Data 7**).

**Figure 5.**
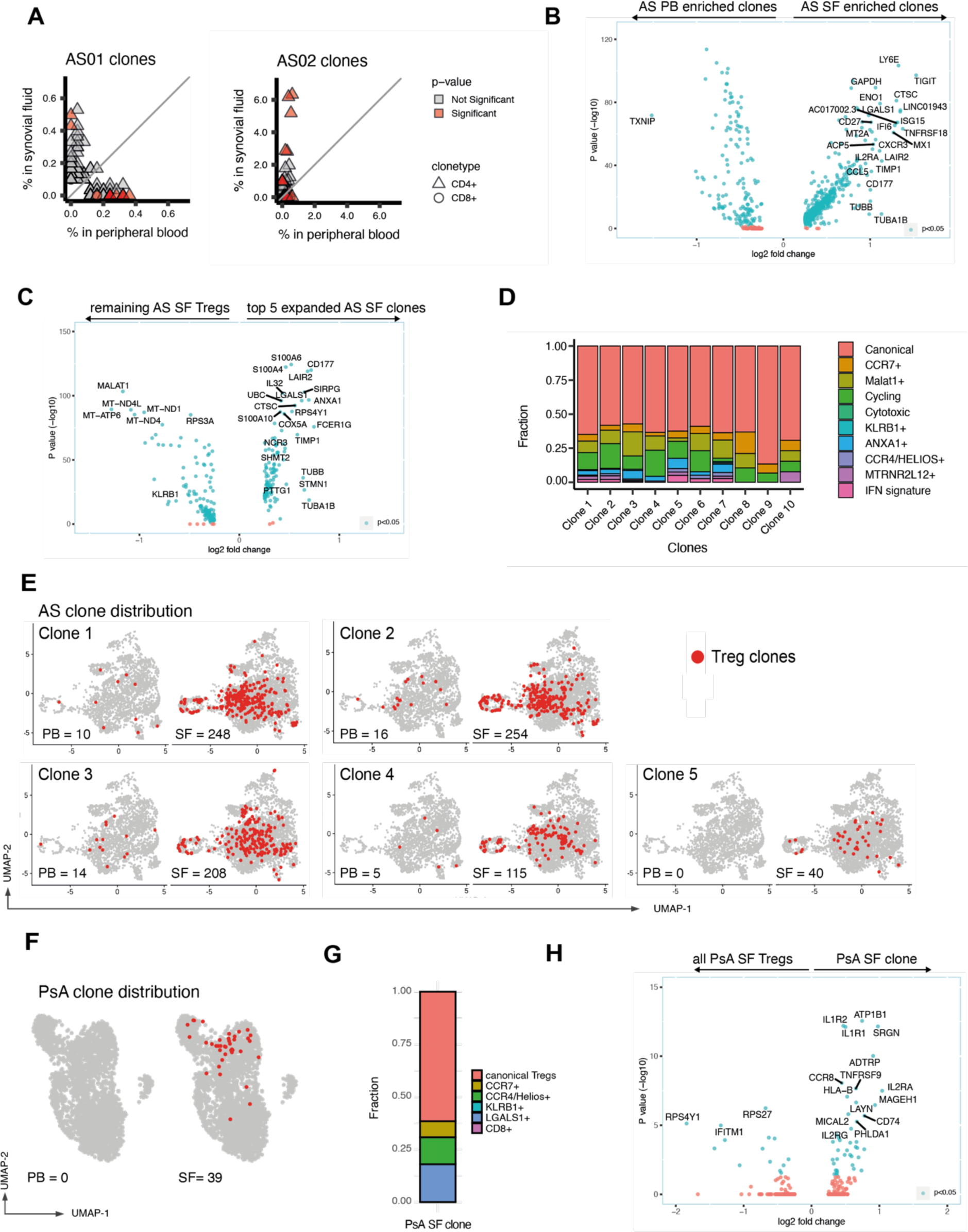
Clonally expanded Tregs with identical TCRα/β sequences and distinct transcriptional features are expanded in the synovial fluid and are present in multiple Treg clusters. **(A)** Individual clonal expansion across PB and SF for two AS patients. Triangles represent CD4 clonotypes. Circles represent CD8 clonotypes (none detected). Data points coloured red show significantly expanded clonotypes (adjusted p value ≦ 0.05). Two-sided Fisher’s exact test, Benjamini–Hochberg correction. **(B)** Differential gene expression of SF-enriched compared with PB-enriched clones. Genes with log fold change >0.25 (and <-0.25) are shown. Genes with adjusted p value <0.05 are coloured in blue. TCR variable chain genes have been removed (full data in **Supplementary Data 7)**. **(C)** Differential gene expression of the top 5 SF enriched clonotypes (ranked by p-value, two-sided Fisher’s exact test, Benjamini–Hochberg correction) compared to the remaining, non-clonal SF Tregs. **(D)** Cluster assignment of Treg clones enriched in AS synovial fluid. **(E)** UMAP plots of AS Tregs showing the top 5 expanded AS clones in red, together with clonal numbers in PB and SF. **(F)** Distribution of the single expanded PsA clone on the PsA UMAP plot. **(G)** Distribution of the PsA clone to each PsA cluster. **(H)** Differential gene expression of PsA clone versus non clonal SF Tregs.

The top 10 clones (by p value) were distributed across several clusters (**Fig. 5D-E**). Whilst the majority were within the canonical Treg cluster, MALAT1+ and Cycling clusters were overrepresented, whereas very little or no clonal presence was observed in the Cytotoxic and in the KLRB1+ cluster. This indicates that sister clones can adopt different phenotypes in the SF (perhaps driven by an antigen). By contrast, the lack of clonal enrichment in the KLRB1+Treg cluster suggests that their activation is not antigen driven (and might be induced by innate inflammatory cytokines or by microbial products, similar to other CD161^-^ cells^12^).

Despite the smaller size of the PsA dataset, we were able to identify the presence of one markedly expanded TCR clonotype in the SF of one patient (**Fig. 5F**). No cells with the identical TCR were found in the PB. On the UMAP plot, the majority of the clone-sharing cells localised close to each other, prevalently in the canonical Treg cluster, although cells were found in other clusters (with the exception of the CD8+ and the KLRB1+ clusters) (**Fig. 5G**). The expanded PsA clone highly expressed *TNFRSF9*, encoding for the costimulatory molecule CD137 (also known as 4-1BB), *CCR8*, *IL1R1* and *IL1R2* (**Fig. 5H**).

These data show that expanded Treg clones are enriched in the SF, and that sister Treg clones selectively enter different Treg clusters to adopt distinct gene expression modules within the inflamed joint.

### A program of genes including the checkpoint inhibitor LAG3 is upregulated by a population of synovial KLRB1+ Tregs

We next wished to further investigate the expression of the checkpoint inhibitor *LAG3*, which we confirmed to be largely confined to the synovial fluid cytotoxic and KLRB1+ subsets in AS (**Fig. 4**), but also in PsA (**Fig. 6A**). Examination of individual Tregs expressing *LAG3* confirmed the increase in the SF (**Fig. 6B**). *LAG3*-expressing Tregs presented an associated suppressive module including the coinhibitory receptor Tim-3 (*HAVCR2*), *IL10*, and its associated transcription factor *MAF* (**Fig. 6C**). We confirmed by FACS that CD161^+^ Tregs express LAG-3 upon activation more frequently than CD161-Tregs (**Fig. 6D-F**). Thus LAG-3 is preferentially expressed on specialised Treg subsets as part of a suppressive module increased in the SF; and LAG-3 protein is upregulated on the surface of AS CD161^+^ Tregs.

**Figure 6.**
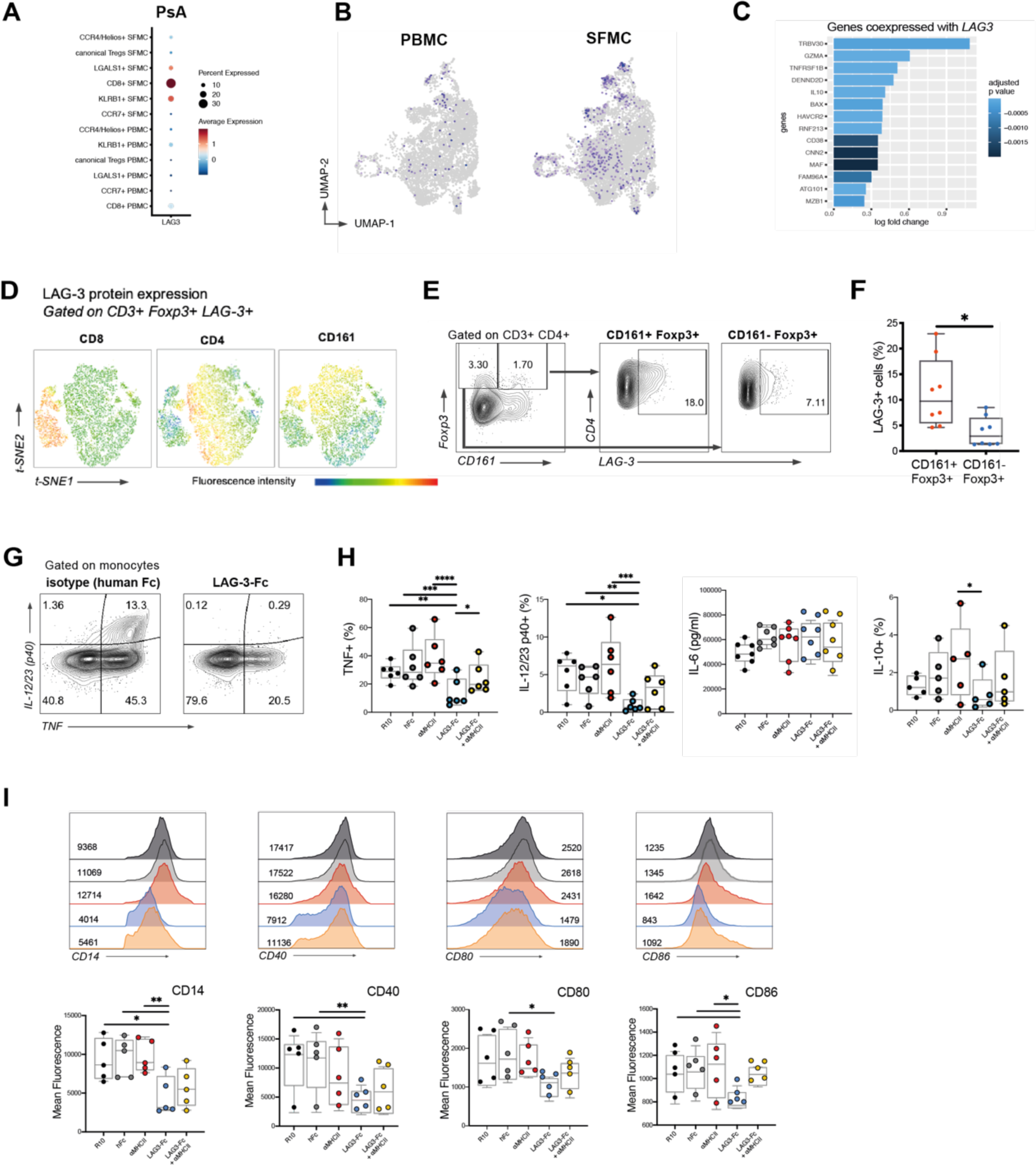
LAG-3 is selectively expressed by CD161 Tregs in SpA and suppresses SpA monocyte TNF and IL-12/23 production and costimulatory molecule expression. **(A)** LAG3-expressing Tregs distribution across different clusters. Bar plot showing fraction of *LAG3*+ cells. (B) *LAG3*-expressing cells distribution over the PB and SF UMAP split plots. **(C)** Genes coexpressed with *LAG3* in SF Tregs: log fold change in gene expression is shown, compared to *LAG3*-cells in SF. **(D)** The majority of LAG-3+ Foxp3+ T cells are found within the CD161 compartment. tSNE plots generated from flow cytometry staining of CD3+ LAG3+ Foxp3+ from one AS patient (after activation with anti-CD3 and-28 for 18 hours), with expression of CD4, CD8 and CD161. **(E)** LAG-3 expression in CD161+ and CD161-Foxp3+ cells, exemplary flow cytometry plot. **(F)** Percentages of LAG-3+ cells in CD161+ versus CD161-Foxp3+ CD4+ cells, and in CD8+ versus CD4+ Foxp3+ cells, from AS PBMCs after activation, n=8. * p<0.05 unpaired non parametric t-test. **(G)** Representative plots of IL-12/23 subunit p40 and TNF expression in CD14+ monocytes in control conditions (left) and in the presence of LAG-3-Fc (right). **(H)** Cytokine production determined by flow cytometric intracellular staining (% of live monocytes), or by concentration in the culture supernatant (for IL-6) after LPS activation in the presence of LAG-3 (and control conditions). **(I)** Costimulatory monocyte surface markers changes after culture with LAG-3 fusion protein. For each marker, a representative stain from one individual (with geometric mean fluorescence intensity for each condition) is shown on top, and data points with minimum and maximum values and IQR on the bottom panels. Boxplots show data points (n=5-7 AS monocytes) with minimum and maximum values and IQR. (* p ≤ 0.05, **≤ 0.01, *** ≤ 0.001, one-way ANOVA).

### LAG-3 suppresses TNF and IL-12/23 production and costimulatory molecule expression by SpA monocytes

We next sought to explore if LAG-3 binding played a functional role in the context of inflammation. We asked if LAG-3 may have actions on inflammatory effector cells. We tested the functional effects of LAG-3 on cytokine secretion from isolated SpA patient-derived CD14^+^ monocytes stimulated with LPS. A LAG-3 fusion protein with a human Fc portion (LAG-3-Fc) consistently inhibited monocyte production of TNF and IL-12/23 p40 (**Fig. 6G-H**). Blockade of HLA-II partially reversed the effect on TNF (**Fig. 6H**).

We next analysed the effect of LAG-3-Fc on the expression of monocyte surface markers. LAG-3-Fc downregulated monocyte CD40, CD80 and CD86 expression, all crucial costimulatory molecules for T cell activation and maturation (**Fig. 6I**). Blocking anti-HLA class II partially reverted the effect, indicating at least a partial competition for the same target. LAG-3-Fc also reduced expression of CD14. Similar results were seen using cultures of whole PBMCs from AS patients, where we observed, with LAG-3-Fc treatment, a downregulation of CD14, CD16, CX3CR1 on CD14^+^ cells, without affecting monocyte viability (**Supplementary Fig. 6**).

In conclusion, LAG-3, is able to restrain monocyte activation and inflammatory cytokine production, at least in part through HLA class II.

## Discussion

This study delineates at the single cell level Treg populations found in the joints of patients with inflammatory spondyloarthritis, identifying both established and novel regulatory subsets. Our analysis, surveying over 16,000 Tregs, represents an atlas of the diverse transcriptional phenotypes acquired by Tregs in the blood and synovial fluid (SF) in the course of spondyloarthritis (graphical summary shown in **Supplementary Fig. 7**). We validate our description of Treg populations independently in two parallel datasets from related human inflammatory diseases, AS and PsA. All Treg subpopulations express a common core set of regulatory transcripts (*FOXP3*, *CD25*, *TNFRSF4*, *TNFRSF18*), with additional specialized modules expressed in a cluster-specific manner. Whilst certain genes, including some with known regulatory function (e.g. *CTLA4* and *TIGIT*) are expressed by multiple Treg subsets, expression of certain functional gene modules is limited to specific Treg populations, a feature further exacerbated in the joints. Thus, *IL10* and *LAG3* are largely (co-)expressed in the KLRB1+ and cytotoxic Treg populations. In the KLRB1+ cluster, we observe expression of the transcription factor ROR-*γ*t, as well as other Th17-associated genes. In view of their lack of expression of *IKZF2,* we propose that cells in this cluster are equivalent to the ROR*γ*t^+^ Helios^-^ Tregs found in the mouse intestine, known key regulators of mucosal immune homeostasis^18^. The analogous gene profile (including expression of *KLRB1*, *CCR6*, *LAG3*, *IL10*, *CTLA4*) has also recently been identified through scRNAseq in the healthy human colon^19^. We and others have proposed that cellular migration between the gut and joint (ie. a “gut-joint axis”) may play a key role in SpA pathogenesis^20^. Our identification of a joint Treg population with strikingly similar gene expression to a colonic Treg population provides evidence (albeit indirect) that specific Treg subsets can also traffic to or between distant inflammation sites.

During inflammation, Tregs can integrate environmental signals to tailor their transcriptional program in a manner that is instrumental for effective regulatory function. In KLRB1+ Tregs, a ROR*γ*t-dependent program could represent a competitional advantage over Th17, or favour colocalisation with Teffs in sites of Th17-driven inflammation^21^. Suppressive markers or transcription factors found enriched in this cluster in the joint have been shown to have regulatory functions in other effector organs, such as LAG-3 in the CNS^22^, and Maf in Th17-driven colitis^23^. Of note, in our data, ROR-*γ*t was the sole T helper-related transcription factor strongly characterizing a Treg subset, the KLRB1+ cluster. *TBX21 (*T-bet) and *GATA3* were in fact not differentially expressed in any of the other Treg clusters. Considering the partially shared differentiation pathway of the two cell lineages, we hypothesize that a transcriptional shift of Tregs towards Th17 in AS and PsA could be instrumental to regulate the heightened Th17 responses present in SpA^24^. Suppressive ability has been in fact demonstrated for CD161^+^ Treg cells obtained from the joints of children with Juvenile Idiopathic arthritis^25^, where they have been described to express gut homing features^26^, and also Crohn’s disease^27^. Of note we, identify *GPR25* as a key gene upregulated in this Treg subset. The function of this (AS-associated) gene is currently unknown but merits further investigation.

In parallel, we observed a population of Tregs expressing molecules of known cytotoxic function (including granzymes). This cluster can be further divided into subclusters expressing either *CD4* or *CD8*. CD8+ Tregs have been identified in autoimmune diseases, tumor immune infiltrates and chronic viral illnesses^28^. Cell contact-dependent mechanisms (e.g. using CTLA-4), tolerization and perforin-mediated cytotoxicity have all been described as contributing to their suppressive function^10^. One study in particular described a very similar phenotype to the one we observed, characterized by markers such as *CCL4* and *LAG3* and a stable expression of *FOXP3*^29^. The chemokines CCL4 and CCL5 (also known as RANTES, among the top upregulated genes in the SF), are chemoattractants for effector T cells that express their ligand CCR5^30^, possibly to regulate their function. Consistently, we previously observed upregulated *CCR5* expression on effector CD8^+^ T cells in PsA SF^15^. The production and release of cytotoxic peptides by Tregs has been described previously: perforin- and granzyme B-expressing Tregs are able to kill autologous antigen presenting cells^31^, and enhanced CD8^+^ cytotoxicity was recently described in the SF of AS patients^32^.

We also observed a synovial Treg signature consistent with exposure to interferons, including *IG15*, *MX1* and *IFI6*. This signature was manifested in an individual subset defined by this phenotype and in global changes seen across all synovial Tregs. Interferons are classically produced in response to viral infections, but might in SpA reflect intracellular bacteria or bacterial products. We also found evidence of TNF exposure: TNF receptor superfamily (TNFRSF) genes were in fact among the top upregulated genes in SF Tregs. Activation of GITR (*TNFRSF18*), OX40 (*TNFRSF4*) and TNF-RII (*TNFRSF1B*) in Tregs, via NF-*κ*B, is known to provide crucial survival and stability signals^33^, and maintain *FOXP3* demethylation^34^. These changes could provide environmental adaptation and survival in a TNF-rich environment such as the SF, while preserving, or even enhancing, their suppressive function. In parallel, expression of these genes may constitute a (non-specific) tissue module or signature, as shown in a recent scRNAseq analysis of murine non lymphoid tissue Tregs^35^.

Use of the 5’ scRNAseq technology here allowed us to study the individual T cell receptor alpha beta pairs of over 16,000 AS and PsA Tregs. We observed selective expansions of individual sister clones in the joints of both AS patients and one of the PsA patients studied (noting the smaller numbers of PsA Tregs may have limited our ability to detect statistically significant expansions). Individual sister clones with identical TCR *α* and *β* nucleotide sequences were detected within different Treg clusters, showing that entry into these populations is not fixed or determined at an early developmental stage and that single clones can have divergent fates. This divergence of fate has been described previously for effector T cells^36^ but to our knowledge not previously been described for Tregs. Moreover, it is also non-stochastic, as evidenced by selective enhancement within specific subsets and the virtual absence of clones within the KLRB1+ subset.

Antigen contact via the T cell receptor (TCR) is an incompletely understood feature of Treg biology, and identification and characterization of antigen-specific Tregs in humans has proven difficult, in part because of their relative rarity in blood. We show that the TCR repertoire of SF Treg cells is diverse and distinct from PB Treg cells, with multiple significant clonal expansions which are unique to the synovial compartment. This suggests that synovial Tregs may have expanded upon antigen contact in the joint or after an encounter in another body site. Significantly expanded clones showed increased expression of markers including *CD177*, a ligand of PECAM1 associated with neutrophil function but recently observed in breast cancer-infiltrating Tregs^37^, and *SIRPG*, whose product mediates adhesion to antigen presenting cell by binding to CD47 potentially optimizing antigen presentation^38^. In the PsA clone we observed upregulation of genes such as *CCR8*, *IL1R1/2*, *MAGEH* and *LAYN*. These markers have been associated to highly suppressive features^39, 40^ or strongly characterizing intratumoral Tregs in two independent studies^41, 42^. Finally *TNFRSF9*, encoding for the costimulatory molecule CD137 (also known as 4-1BB), has been reported to identify antigen-specific Tregs in immune driven conditions^43^.

Intrigued by the selective and marked upregulation of *LAG3* expression in two SF Treg clusters, we explored its functional role in modulating inflammatory responses. LAG-3 (Lymphocyte activation gene-3, also known as CD223), is a transmembrane protein which shares 20% homology at the amino acid level with CD4^44^, and it binds a common ligand, the HLA class II molecule. Expressed on a number of T cell subsets (including Tregs), because of its expression by tumor infiltrating lymphocytes^45^ often in association with exhaustion markers, LAG-3 has become an important potential target for immunotherapy^46^. While being studied as a new possible checkpoint inhibitor to treat solid tumors, its role and its mechanism of action on Tregs are not clear. LAG-3 is expressed by Tregs upon activation, and conditional *Lag3* knockout Tregs exhibit reduced functionality^47^, suggesting it might confer suppressive activity. While the first reports studying LAG-3 binding to myeloid cells observed dendritic cell (DC) activation^48^, one following report^49^ described downregulation of CD86 on bone marrow derived DCs following MHC class II binding by LAG-3-expressing Tregs. LAG-3 on Tregs was also highlighted in an innate like cell (ILC)3-driven experimental inflammatory model of colitis, where LAG-3^+^ Tregs specifically targeted CX3CR1^+^ macrophages, decreasing their production of IL-23 and IL-1*β*, ameliorating disease^50^. Interestingly, CX3CR1^+^ macrophages are increased in many inflamed tissues in AS, including the intestine^51^. Together, these lines of evidence indicate that LAG-3 might represent a suppressive mechanism in gut Tregs, which have the potential to traffic to other sites, perhaps in response to Th17-driven inflammation. Our description of enhanced LAG-3 expression in SpA SF (Tregs) is we believe the first in human inflammatory arthritis.

Our data confirm the co-expression of *LAG3* and *IL10*, suggesting they are part of a coordinated suppressive programme specific for tissue immunity. Interestingly, IL-27 has been reported to promote simultaneous LAG-3 expression and IL-10 production in murine and human T cells^52, 53^. Supporting a potential application of the LAG-3-mediated suppression in inflammatory arthritis, its natural target, the HLA class II complex, is highly expressed on inflammatory synovial monocytes^54^, DCs^55^ and macrophages^56^. Whilst we have not directly confirmed the inhibitory capacity of LAG-3^+^ Tregs, we have shown that LAG-3 inhibits production of TNF and IL-12/23 and induces downregulation of activation markers CD40, CD80, CD86 on myeloid cells. TNF and IL-12/23 are key inflammatory cytokines whose inhibition has been shown to be of therapeutic efficacy in SpA, hence our data support a novel LAG-3-based therapeutic approach for SpA and related inflammatory mediated diseases. The inhibitory mechanism we describe could additionally have a role in limiting Th17 responses.

Although we do not know if these populations are specific to synovial Tregs (or indeed SpA joints), we identify specialized subsets including one expressing *KLRB1* and *LAG3* (which we show can suppress SpA monocyte inflammatory responses) akin to an intestinal mouse Treg subtype, and a cytotoxic Treg population that includes a significant CD8^+^ regulatory subset. Another limitation of our study is the limited number of patients studied with scRNAseq. We mitigated this by studying a large number of cells and confirmed our findings across two independent datasets and additionally verified the key findings using FACS analysis of additional patients.

In conclusion, we here present a large human Treg data set in the context of inflammation, that shows distinct Treg subsets and identifies a broad transcriptional profile upregulated across all synovial Tregs. TCR analysis shows that sister clones can specifically enter different subsets and provides evidence of Treg clonal expansion which may be driven by antigen. Our in-depth characterization of Treg subsets shows specific and coordinated expression of LAG3 on certain Treg subsets. Demonstration of LAG-3 function allows us to identify potential novel therapeutic approaches both for autoimmune diseases (mimicking Treg functions) or malignancy (by inhibiting Treg functions).

## Methods

### Participant recruitment, ethical approval

Patients with AS and PsA were recruited during routine clinical care following written informed consent in accordance with the protocol approved by the South Central – Oxford C Research Ethics Committee (IFIA, Immune Function in Inflammatory Arthritis: ethics reference 06/Q1606/139). All patients (**Supplementary Table 1**) fulfilled the disease classification criteria (respectively ASAS and CASPAR)^57, 58^, and were naïve to biologic disease-modifying antirheumatic drugs (DMARD) and not on any conventional DMARD at the time of the sample. All patients with AS were HLA-B27 positive with evidence of active axial and peripheral joint involvement. Patients with PsA had large joint peripheral oligoarthritis although none were HLA-B27 positive. Synovial fluid samples were obtained during knee joint aspiration performed for therapeutic reasons.

### Cell isolation and flow cytometry

SFMC and PBMC were freshly isolated within 30 minutes of sample collection by density-gradient centrifugation using Histopaque (Sigma). For flow cytometric analysis, samples were prepared by washing cells twice using FACS buffer (Phosphate-buffered saline (PBS) with the addition of 1% Fetal Bovine Serum (FBS)) in 96 well U-bottom plates (Corning) or round-bottom polystyrene tubes (BD Biosciences). 0.2-0.5 x 10^6^ cells per well or tube were stained. Staining buffer was prepared by adding flourochrome-conjugated antibodies and fixable dyes (list in **Supplementary Table 2**) to FACS buffer and used to resuspend cell pellets for staining mixing thoroughly. Cells were then incubated for 20 minutes at 4°C in the dark. Where surface staining included an antibody for LAG-3, cells were cultured overnight with plate-bound anti-CD3 (OKT3, 1 μg/ml, Biolegend) and soluble anti-CD28 (CD28.2, 1 μg/ml, Thermofisher), followed by cell staining at 37°C. After staining, cells were washed twice with 200 μl FACS buffer and resuspended in 200 μl fixing buffer (PBS with the addition of 3% paraformaldehyde) before acquisition. When staining for intracellular proteins, after completing the staining for surface markers as described above, cells were first permeabilised by resuspending them in a fixation/permeabilization solution (Cytofix/Cytoperm, BD Biosciences) at room temperature for 30 minutes, then washed twice in 200 μl Permwash buffer (BD Biosciences), and finally stained for intracellular proteins before being suspended in fixing buffer for the acquisition. For the detection of intracellular cytokines in monocytes, Brefeldin A (GolgiPlug, BD Biosciences) was added before staining. When staining T cells for Foxp3 or Helios, a variation of the intracellular staining protocol was adopted, using, instead of Cytofix/Cytoperm and Permwash, the equivalent products in the FOXP3 / TF Staining Buffer Set (Thermofisher). Sample acquisition was performed on a BD LSR Fortessa flow cytometer. Calibration and setup was performed daily with BD FACSDiva CS&T Beads (eBioscience). Compensation was performed using single stained One Comp eBeads (eBioscence), or for the Viability dye, using primary cells. Results were analysed using FlowJo software (v. 10.6.2, Treestar). Dimensionality reduction of flow cytometry data and t-SNE plot generation was obtained using the “t-SNE” Flowjo plugin.

### Fluorescence activated cell sorting (FACS) for scRNAseq

After isolation by density centrifugation, PBMC and SFMC were immediately stained with fluorescently conjugated antibodies in RNAse-free PBS, 2 mM EDTA and then FACS-sorted prior to droplet-based single-cell RNA sequencing. AS samples were stained with the following antibodies: CD3-PerCP-Cy5.5 (OKT3), CD8a-PE (RPA-T4), CD45RA-PE/Dazzle (HI100), CD25-PE (BC96), CD127-PE/Cy7 (A019D5) (all from Biolegend, and used at 1:50 dilution) and Fixable Viability Dye eFluor520 (eBioscience, dilution 1:250) to exclude dead cells. Cells were sorted on a Sony SH800Z. Memory Tregs were sorted as in **Supplementary Fig. 1A** (CD45RA^-^ (negative) CD3^+^ CD25^+^ CD127^low^). Cells were then collected in a collection buffer (Phenol Red-ve RPMI + 4% Bovine Serum Albumin + Hepes 25mM). After sorting, cells were stained separately with Fc blocker (concentration 1:20) (TruStain FcX, Biolegend), rested for 15 minutes, then washed, then resuspended in buffer to be further stained with with the oligo-tagged TotalSeq™-C0251 Hashtag antibody. Cells were again washed twice with FACS buffer then kept on ice until loaded onto the Chromium controller. For sample AS02, PBMC and SFMC were not processed fresh but thawed after being cryopreserved in liquid nitrogen. For PsA samples, T cells were sorted and prepared for sequencing as previously described^15^.

### 10x Genomics single cell RNA library preparation

Cells were counted and loaded into the chromium controller (10x Genomics) chip following the standard protocol for the chromium single cell 5′ Kit (10x Genomics). The total time taken from sample retrieval to upload on the chromium chip was 4 h. A cell concentration was used to obtain an expected number of captured cells, approximately 15,000 cells per sample. All subsequent steps were performed based on the standard manufacturer’s protocol. Libraries were pooled and sequenced across multiple Illumina HiSeq 4000 lanes to obtain a read depth of approximately 30,000 – 40,000 reads per cell for PB and SF gene-expression libraries of both patients, and 6,000 or 20,000 reads per cell for V(D)J-enriched T-cell libraries from both PB and SF for patients AS01 and AS02 respectively (**Supplementary Table 3**). Chromium 10x V(D)J single-cell sequencing data were mapped and quantified using the software package CellRanger (v2.1 for the PsA samples and v3.1 for the AS samples) against the GRCh38 reference provided by 10x Genomics with that release.

Demultiplexing of the Illumina files and generation of fastq files containing the scRNA-seq data were performed using the “*mkfastq*” function in the Cell Ranger software package. Alignment of scRNA-seq reads to the human reference genome (GRCh38) and transcript quantification were performed using the “*count*” function in Cell Ranger (10x Genomics, v. 2.1 for PsA samples, and v. 3.0.2 for AS samples). All five samples had similar coverage in terms of unique mRNA molecules and genes represented, altogether surveying a total of 33,694 genes (sequencing metrics detailed in **Supplementary Table 3)**. The generated consensus annotation files for each patient and sample type (blood or SF) were then used to construct clonality tables and input files for further downstream analysis using the jsonlite (v. 1.6.1) package

### Single cell RNAseq analysis

#### Quality control

Downstream analysis of the count matrices was carried out using R (version 3.6.1) and the Seurat package (v. 3.1.4). After cell-containing droplets were identified, gene-expression matrices were first filtered to remove cells having >10% mitochondrial gene transcripts, <250 or >4,000 genes expressed or >25,000 UMI (Unique Molecular Identifiers). The Seurat demultiplexing function (“*HTODemux”*, with a threshold set at the 99^th^ quantile of the negative binomial distribution for the oligo) was then used to demultiplex the hashing library in order to identify Tregs and to remove doublets. Cells were further filtered to exclude cells not expressing any transcripts from CD3 complex-associated genes (*CD3E*, *CD3D*, *CD3G*) and TCR multiplets (defined as cells with greater than 1 TCR beta chain or greater than 2 TCR alpha chains).

To further remove any CD14+ cells or multiplets which may have escaped exclusion by cell sorting, a preliminary round of dataset integration, dimensionality reduction and cell clustering as described below was used to identify cells belonging to CD14+ clusters. These cells, along with any additional cells expressing CD14, were then excluded from the input used in generating a final integrated dataset.

Quality control of the PsA dataset is detailed elsewhere ^15^: briefly, similarly to the AS dataset, cells with >10% mitochondrial gene transcripts; <500 or >3,500 genes; >25,000 UMI; expressing both *CD4* and *CD8;* or with greater than 1 TCR beta chain or greater than 2 TCR alpha chains were removed.

### Dataset integration

The analytical strategy used to integrate data from different tissue samples and across different experiments uses the Seurat v.3 pipeline ^59^. QC-filtered matrices from all patients were individually normalised using the “SCTransform” function before running the “SelectIntegrationFeatures” function to determine the top 3000 variable genes. TCR genes were then excluded from these variable features and the matrices integrated according to the standard Seurat version 3 SCT integration pipeline (“PrepSCTIntegration”, “FindIntegrationAnchors”, “IntegrateData”). TCR genes were excluded from variable features to prevent downstream clustering based on clonality, which can differ between patient samples and potentially distort clustering based on cell phenotype. A Treg cluster that had previously been identified within a PsA CD4/CD8 10x dataset analysed by our lab^15^, was exported as a raw object using Seurat’s “*SubsetData”* command for comparison with AS Tregs. These PsA Treg cells were re-clustered and re-analyzed applying the same pipeline detailed above, set to default parameters.

An integrated dataset consisting of Tregs from both the AS and PsA datasets was also created following the same methods outlined above, additionally regressing out the number of UMIs and percentage of mitochondrial transcripts (vars.to.regress argument of the SCTransform argument).

### Dimensionality reduction and clustering

The function “*RunPCA*” was performed on the integrated assay to compute principal components (PC), the first 30 of which were selected, based on the Seurat elbow plot, and specified as the dims argument to the “FindNeighbors” and “RunUMAP” functions. Clusters were then discovered by the “*FindClusters*” function at a resolution of 0.2 according to the standard Seurat workflow. Each cluster was classified by differentially expressed genes and visualised by a Uniform Manifold Approximation and Projection (UMAP) plot. The same steps, consisting of finding principal components, construction of an SNN graph and clustering, were applied to the PsA gene expression matrices, specifying a resolution of 0.3 to the Findclusters function.

### Differential gene expression analysis

Pairwise differential gene expression comparisons were made across cell clusters or conditions. Differential expression analysis was performed using negative binomial generalized linear model implemented in Seurat, through the command “*FindMarkers*” or “*FindAllMarkers*”, considering markers expressed in at least 10% of cells. Wilcoxon rank-sum test with Bonferroni correction was used to determine differences. To perform statistical analysis of functional profiles the R package Clusterprofiler (v. 3.14.3) was used. Genes with significant differential expression for each pairwise comparison were used as input for the pathway analysis, using the Gene Ontology or the Reactome Pathway^60^ repositories. The gene module score was calculated with the command “*AddModuleScore*” in Seurat^59^.

### T cell receptor reconstruction and analysis of clonality

Chromium 10x V(D)J single-cell sequencing data were mapped and quantified using the software package CellRanger (v2.1 for the PsA samples and v3.1 for the AS samples) against the GRCh38 reference provided by 10x Genomics with that release. The generated consensus annotation files were then used to construct clonality tables and input files for further downstream analysis. After TCR reconstruction, the proportion of cells having the same clone was compared between sample types for each clone using a two-sided Fisher’s exact test with Benjamini and Hochberg correction for multiple comparisons (R Stats Package) considering all clones with 3 or more cells in either synovial fluid or peripheral blood. Clonotypes were defined as cells having identical complementary determining region 3 (CDR3) nucleotide sequences for the alpha and beta chain CDR3 sequences assigned to each cell. As it was not possible to deduce beta- and alpha-chain pairing for partitions with multiple beta chains, these partitions were treated as a single clone. When analysing both gene expression and clonality of the same cells, partitions containing more than one beta chain or more than two alpha chains were considered multiplets and were excluded from analysis.

### Monocyte LPS stimulation

Monocytes were isolated using a CD14+ magnetic positive selection kit (CD14 Microbeads, Miltenyi Biotec) from patients PBMCs, achieving a purity of 85-95%. Isolated CD14+ cells (or, in some experiments, whole PBMCs) were plated at a concentration of 0.5 x 10^6^ cells per well in 96-well round-bottom plate. LPS (LPS-EB, Invivogen) was added at a dose 10 ng/ml. After LPS stimulation, cells were kept in culture overnight. When determination of intracellular cytokine production was desired, brefeldin A was added four hours after LPS stimulation. For experiments that evaluated the effect of LAG-3 ligation on monocytes, a recombinant human LAG-3 IgG1 Fc chimera protein (R&D) at a concentration of 2.5 μg/ml was used. Recombinant human IgG1 Fc control and anti-human HLA class II (anti-DQ, -DR-DP) (clone Tu39, BD Biosciences) were also used at 2.5 μg/ml, and were added 2 hours before the addition of LPS.

### Statistical analysis

Statistical analysis was performed using the software GraphPad Prism 8.4. Data are presented in the form of box-and-whisker plots (minimum, maximum and interquartile range). Statistical analysis on the scRNA-seq data was performed using the R “Stats” package or the built-in statistical tools for each R package used.

## Supplementary Information

Supplementary Figure 1. Treg sorting strategy, patient contribution to Treg clusters and expression of cluster-characterizing genes.

Supplementary Figure 2. Pathway analysis of genes enriched in Treg clusters confirms an interferon-responsive signature cluster; co-expression of key co-inhibitory lineage and transcription factors.

Supplementary Figure 3. Th17-like Tregs are more often TIGIT^low^ compared to other Tregs, and C8+ Tregs express *FOXP3* and *IL2RA* at levels comparable to CD4 Tregs.

Supplementary Figure 4. Single cell analysis of PsA Tregs confirms cluster identities found in AS.

Supplementary Figure 5. Tregs are increased in frequency in SpA synovial fluid and show specific gene expression changes.

Supplementary Figure 6. Soluble LAG-3 protein modifies monocyte phenotype downregulating selected activation and maturation markers.

Supplementary Figure 7. Graphical summary.

Supplementary Table 1. Demographic and clinical data of patients studied with scRNAseq

Supplementary Table 2. Flow cytometry antibodies used

Supplementary Table 3. scRNAseq runs and sequencing metrics

Supplementary Table 4. Gene modules list

Supplementary Data 1. AS Differentially enriched genes by cluster

Supplementary Data 2. AS Synovial fluid differentially enriched genes

Supplementary Data 3. Pairwise gene expression comparisons

Supplementary Data 4. SpA (integrated AS/PsA) differentially enriched genes by cluster

Supplementary Data 5. Enriched AS clones

Supplementary Data 6. Enriched PsA clone

Supplementary Data 7. Differentially expressed genes in top 5 SF clonotypes vs non clonal SF

## Acknowledgments

We thank the patients for their participation

## Author contributions

DS, FP, HAM and PB conceived and designed the experiments; DS, FP, and HAM performed the 10× experiments, designed and performed the computational analysis aided by SS; AR assisted with the cell sorting; FP and HAM generated the PsA dataset, DS performed the flow cytometry and the in vitro experiments; SS and AR contributed to the discussions; DS, FP, HAM and PB wrote the paper; HAM and PB co-directed this study. All authors read and approved the paper.

## Data and materials availability

Raw expression matrices will be deposited on GEO. The code used in data analysis and for the generation of figures will be made available anytime upon request or at final submission.

